# Diet alters epidemic size and timing in a trophically-transmitted parasite

**DOI:** 10.64898/2026.05.15.725575

**Authors:** Juliana Jiranek, Allyson Motter, Neha Channamraju, Ean Huang, Tessa Batterton, Amanda Gibson

## Abstract

A host’s diet can alter the course of parasite infection. This is especially true of trophic parasites, which a host acquires through feeding. While a large body of work attests to the role of diet in the spread of disease within-hosts, diet can also impact host density and encounter rate with parasites, both of which are expected to modify disease dynamics. When parasites are acquired through feeding, epidemics may be larger and more severe on high-quality diets if these diets support a higher density of hosts that feed more and thus ingest more parasites. Alternately, epidemics may be more severe on low-quality diets if malnourishment decreases hosts’ ability to resist disease. To differentiate these hypothesized effects of diet on disease, we quantified individual infections and epidemic dynamics for the natural microsporidian parasite *Nematocida ironsii* infecting its nematode host *Caenorhabditis elegans*. We measured feeding rate, parasite transmission, and host fitness across three bacterial diets that vary in quality and elicit distinct feeding behaviors in *C. elegans*. We found that low-quality diets reduced feeding rate, which corresponded to reduced acquisition of parasite spores. However, these diet-mediated differences in parasite acquisition did not directly map onto fitness consequences: hosts eating the poor-quality diet had similar reductions in fitness to those on higher quality diets. During epidemics, a combination of increased parasite acquisition and higher population growth rates resulted in higher parasite abundance for hosts on high-quality diets. Our work underscores the importance of considering both individual- and population-level impacts acting in concert to determine how diet affects the spread of infectious disease.

## Introduction

Hosts commonly encounter parasites while they are eating. As a result, diet can have a large impact on a host’s probability of contracting an infectious disease and spreading it to other hosts (Nunn and Altizer 2006, Adler et al. 2018). Variation in diet quality may generate heterogeneity in the spread of disease, which makes it more challenging to predict and manage epidemics (Smith 2007). This problem is exacerbated by the fact that dietary quality may alter multiple host traits simultaneously, making it difficult to determine which ones are most important in driving epidemic patterns (Cressler et al. 2014, Hite and Cressler 2018, Pike et al. 2019). To better understand how dietary quality impacts epidemic size and timing, we explored the contribution of multiple host traits to parasite transmission across scales.

Most simply, a host’s diet dictates foraging rate, which determines encounter rates with trophically-transmitted parasites. For example, zooplankton hosts on high-quality diets ingested more fungal spores because they ate more (Penczykowski et al. 2014). Alternatively, individuals on low-quality diets may increase their foraging rate to meet nutritional demands, thereby increasing parasite exposure (Cruz-Rivera and Hay 2000, Fink and Von Elert 2006, Wang et al. 2006). Higher intake of parasites could increase the probability that the parasite establishes and could result in a larger infections overall (Hall et al. 2012).

Additionally, diet quality impacts parasite growth within a host. On one hand, increasing diet quality could increase host immune function (Moret and Schmid-Hempel 2000; Kraaijeveld & Godfray, 1997; Fellous and Lazzaro 2010), leading to a decrease in disease and/or its fitness consequences (Hite and Cressler 2018). Alternatively, increasing dietary quality could increase pathogen growth because resources are more abundant within the host (Bedhomme et al. 2004) or because the host is better able to tolerate the infection (Sternberg et al. 2012). Because hosts with larger infections tend to shed more parasites to their environment and/or fellow hosts (de Roode et al. 2009), this variation in within-host spread can shape epidemics.

Dietary quality could also modulate disease transmission through impacts on host population size and density. Basic consumer–resource models predict that higher quality diets support larger populations (Griffen and Drake 2008). Foundational disease theory suggests that density-dependent diseases spread faster in large, dense populations (Anderson and May 1981). The integration of these two lines of theory predicts that hosts on higher quality diets should experience larger epidemics (Smith 2007). Consistent with this idea, studies in a zooplankton–fungus system suggested that increased host birth rates increased the number of susceptible individuals in the population, leading to larger epidemics (Hall et al. 2009a, Civitello et al. 2015). These findings motivate experimental investigations to directly link diet to transmission.

Building on this prior work, we explored how the effects of diet on disease scale from the individual hosts to epidemics in growing host populations. We used the tractable system of the bacterivore nematode host *Caenorhabditis elegans* and its natural microsporidian parasite, *Nematocida ironsii*. These hosts ingest parasite spores during feeding, then shed spores after a systemic infection establishes in the gut (Troemel et al. 2008). Because the generation time of host and parasite occur on the order of days, we can monitor both individual-and population-level progression of disease. We tested how host diet influenced disease dynamics using three distinct strains of *Escherichia coli* that represent a gradient in dietary quality and host preference (Shtonda and Avery 2006, You et al. 2008a, Brooks et al. 2009, So et al. 2011b, Jiranek and Gibson 2023). To test for differences in parasite acquisition, we measured foraging rates and spore uptake on different diets. We then measured parasite load and host fitness following controlled inoculations to test the effects of diet on a host’s ability to resist disease on different diets. Finally, to evaluate the impact on epidemics, we monitored population growth and disease spread in growing host populations. Our results indicate that dietary quality mediates the size and timing of epidemics through effects on parasite acquisition and population growth rate.

## Methods

We used experimental populations of the nematode *C. elegans* and its microsporidian parasite *N. ironsii* to test the effect of host diet on epidemics. First, we measured host feeding behavior and assessed how this correlated with parasite acquisition. Then, we conducted a fecundity assay and population growth assay to measure the fitness consequences of parasite exposure. Finally, we monitored the progression of experimental epidemics to quantify the dynamics of disease spread (Fig. S1).

### Study System, Strains, and Culturing

*Caenorhabditis elegans* is a model nematode that feeds on transient patches of bacteria in the wild (Félix and Braendle 2010). *Nematocida* species are natural, trophically-transmitted microsporidian parasites of *Caenorhabditis* (Troemel et al. 2008, Zhang et al. 2016). Hosts acquire spores from the environment while foraging, and these spores establish in host intestinal cells (Troemel et al. 2008). It takes about two days (∼52 hours at 20°C) for hosts to begin shedding new spores (Balla et al. 2016). We selected the N2 line of *C. elegans* as our host because of its high baseline susceptibility to *Nematocida*, and we used *Nematocida ironsii* as the parasite (ERTm5 strain; Tecle & Troemel, 2022). Nematodes were kept at 20°C during all experiments.

Stocks of *N. ironsii* spores were generated by infecting large populations of *C. elegans,* isolating spores from the environment and infected hosts after several days of proliferation, then freezing aliquots at −80°C after ensuring spores were free of contaminants (Bubrig et al. 2022; Bakowski et al. 2014). With the exception of the Epidemic Progression assay (see below), the spore dose for assays was 4.45×10^6^ spores per 60-mm exposure plate, which is comparable to doses used in prior studies (Szumowski *et al*., 2014; Balla *et al*., 2015; Bubrig et al. 2022). We used a lower dose of 2.59 x 10^6^ spores per plate for the Epidemic Progression assay because of its longer duration. Because *N. ironsii* spores were harvested from suspensions of lysed infected hosts, all control conditions received a control inoculation of equal volume derived from suspensions of lysed healthy worms. Spores from a single stock were used in all experiments.

To simulate variable diets, we fed *C. elegans* three different strains of the standard bacterial food *Escherichia coli*: DA837, OP50, and HB101.These strains vary in their quality and their palatability to *C. elegans* (Shtonda and Avery 2006, You et al. 2008, Brooks et al. 2009).

OP50 is the standard laboratory food for *C. elegans* and is of intermediate quality (Shtonda and Avery 2006). DA837 is derived from OP50 but has smaller cells that clump together, making them more difficult to ingest (You et al., 2008). These small cells also have less protein and fatty acids by weight compared to OP50 and HB101 (Brooks et al. 2009). *Caenorhabditis elegans* avoid it in favor of other bacteria in choice tests and spend more time roaming and less time eating when they are fed it (Shtonda and Avery 2006). HB101 is a hybrid of *E. coli* strains B and K12 that has a higher mass of carbohydrates and protein per cell than OP50 or DA837 (Brooks et al. 2009). A diet of HB101 increases host body size, satiety responses, and population growth rate (You et al. 2008, So et al. 2011b, Jiranek and Gibson 2023).

Bacterial cultures were grown from frozen stocks in lysogeny broth (LB; Fisher Scientific BP9723-500) inoculated with a single colony at 28°C. Nematode growth medium (NGM; Fisher Scientific NC9379170) plates were seeded with bacteria at a standardized concentration of 5×10^7^ CFU/ml and incubated overnight at 28°C. For the assays of longer duration (Population Growth and Epidemic Progression), we added 1.5% agarose to the NGM to make the medium resistant to digging. For the Epidemic Progression assay, denser cultures (6.7×10^7^ CFU/ml) were used to seed the plates and lawns were grown over a longer period to sustain larger host populations.

### Parasite contact: Bead Acquisition Assay

To determine whether differential foraging could alter parasite contact rates, we first tested if hosts fed at different rates across diet conditions. In accordance with past work (Troemel et al. 2008), we found that *C. elegans* do not change their feeding rate in response to *Nematocida* (Supplementary Materials, Appendix A), so we carried out trials on unexposed plates. Hosts were age-synchronized by isolating eggs using a standard bleach wash (Porta-de-la-Riva et al., 2012). Eggs were added to DA837-, OP50-, and HB101-seeded 100 mm plates. Upon reaching the L1 stage 1,000 hosts were then moved to 60 mm plates seeded with the same bacterial strains mixed with 1.27×10^10^ 0.10 micron fluorescent polystyrene beads (Polysciences; Kiyama *et al*., 2012). After thirty minutes of feeding, hosts were washed off the plates, and the suspension was run through a 10 μm diameter cell strainer to remove hosts from excess bacteria and beads. Hosts were anesthetized with 20 mM sodium azide and mounted on agar pads on microscope slides. We photographed slides at 10X magnification under a compound fluorescent microscope using brightfield and Texas Red filters (Leica DM6B). We used ImageJ to quantify the area of red fluorescence in the host’s intestinal lumen (number of hosts = 170; DA837 = 49 hosts across 7 plates, OP50 = 78 hosts across 8 plates, HB101 = 43 hosts across 7 plates).

The data from the feeding assay were zero-inflated and followed a negative binomial error distribution. To account for this, we fit zero-inflated models with the “glmmTMB” package and a negative binomial (“nbinom1”) distribution (Brooks et al. 2017). We used these models to test the effect of diet on the area of a host’s gut filled with fluorescent beads, with the plate the host foraged on as a random effect. We expected foraging, and thereby bead acquisition, to increase with host size. To determine whether we could use host size as a covariate, we first tested for an interaction effect between host size and diet on bead ingestion. Because we did not detect an interaction, we removed the interaction term and tested the effect of food type with host size as a covariate. We evaluated the impact of model terms using a Wald χ^2^ test. We then performed post-hoc Tukey tests for all pairwise comparisons using the “emmeans” package (Lenth 2023). We also tested the effect of diet on host size. Because the size data failed to meet ANOVA assumptions, we performed a non-parametric Kruskal-Wallis test and a post-hoc pairwise Wilcoxon rank sum test to assess pairwise differences between diets.

### Parasite contact: Parasite Acquisition Assay

To determine if bacterial diet affects parasite acquisition, we measured parasite load after a 24-hour exposure to *N. ironsii*. We added synchronized L1 hosts to DA837-, OP50-, and HB101-seeded 60 mm plates with spores. After 24 hours, hosts were washed off the plates and fixed in acetone. *Nematocida ironsii* was tagged using a fluorescence in-situ hybridization (FISH) probe that binds to *Nematocida* ribosomal RNA in germinated sporoplasms (Troemel et al. 2008), and then hosts were mounted on agar pads on microscope slides. Slides were photographed at 10X magnification under a compound fluorescent microscope using the Texas Red filter. To quantify parasite load, we cast images in black and white in ImageJ, heightened the contrast, and manually counted sporoplasms per individual. These sporoplasms indicate spores that have successfully germinated and established in the host gut. The sporoplasm count approximated a normal distribution when log-transformed, so we tested hypotheses using a linear model. To test if host size could be used in the model as a covariate, we tested the interaction of diet and host size on the log-transformed sporoplasms count. The interaction was not significant, so the final model tested the effect of host size and diet on log-transformed sporoplasm count.

### Fitness: Fecundity Assay

We measured the fecundity of exposed and control hosts on each diet to assess whether diet altered the fitness consequences of disease. At the L1 stage, 500 hosts were either exposed to 60 μL of spore solution or equal volume of a control solution for 24 hours. At 24 hours, when hosts had reached the fourth larval (L4) stage, we transferred individuals to 35 mm plates seeded with the same bacteria as their original plate (n = 130; 16-28 hosts per food×exposure combination). We moved each host to a new plate every 24 hours for six days, during which hosts laid eggs. Plates with eggs were incubated at 20℃ for 24 hours, after which we counted live offspring from each host plate. We censored data from hosts that were lost due to error during transfer or burrowing into agar (n = 9).

We evaluated both daily and lifetime offspring production. We fit a linear mixed model to lifetime fecundity, including fixed effects of diet, parasite exposure, and their interaction. We also fit fixed effects for the block of the experiment and the experimenter who moved the host; this approach is more conservative than fitting random effects given that these factors had fewer than 5 levels (Gomes 2022). Because earlier reproduction increases population growth rate (Hodgkin and Barnes 1991), we counted the offspring produced each day. We modeled the effect of diet, exposure, day, and their 3-way interaction on daily fecundity. We used the same effects as in the lifetime fecundity model but also included a random effect of host ID because of repeated measures. Because many hosts made no offspring in later days of reproduction, we implemented a hurdle model in “glmmTMB” with a negative binomial (“nbinom1”) error distribution and a zero-inflation term that took the day of the experiment into account.

### Fitness: Population Growth Assay

To test the effect of diet on population growth, we compared the size of populations exposed to parasites to that of control (non-exposed) populations after six days of expansion. Following methods described in Jiranek et. al (2023), we reared hosts on DA837-, OP50-, or HB101-seeded 100 mm plates and then either mock-exposed or exposed hosts for 24 hours on the same diet. Then, we picked control and exposed L4 hosts individually onto 100 mm plates of the corresponding diet (n = 60, with one host per 100mm plate; n = 20 for each diet x exposure combination). Five populations were removed from the analysis because the original host did not produce offspring. Four of these populations were in the control treatment, and all diets were represented. Thus, lack of offspring was unlikely to be a treatment effect and their exclusion should not bias our analysis. We allowed the remaining populations to expand for six days (∼2-3 generations) before washing them off their plates and suspending them in a volume of 14.5 ml. To estimate population size, we counted the number of hosts in six 20 μl aliquots per population and multiplied by the average count by the conversion factor 725 (hosts per 20 μl × 725 = hosts per 14.5 mL). We used a generalized linear model with a negative binomial error distribution to estimate the effects of diet, exposure, and their interaction on population size.

### Population: Epidemic Progression Assay

To assess how diet may alter the progression and size of an epidemic, we compared parasite prevalence and population size at four time points throughout the course of an epidemic. We added 50 synchronized L1 hosts to 100mm plates seeded with either DA837, OP50, or HB101 (n = 60) and 35 μl of spores. Populations were allowed to grow for two days, at which point we began to destructively sample five populations per bacterial strain per day for four successive days (n = 5 per diet×day combination). Populations were diluted in a known volume of M9, such that total population size could be estimated from six 20 μl aliquots as in the Population Growth assay.

To assess parasite prevalence and burden, we preserved populations in acetone and treated them with a fluorescent probe, as in the Parasite Acquisition assay. We took images of 50 hosts per population. Fluorescence was quantified in ImageJ. Briefly, we cast images in black and white and set a threshold brightness to select the outline of an individual host, measuring the area of the host’s body. Then, we reset the threshold brightness to only highlight *N. ironsii* fluorescence. We used the “Summarize” function to calculate the total area of fluorescence and the percentage of the host’s body that was fluorescing. If the area occupied by *N. ironsii* was greater than zero, we considered a host infected. We used the number of infected individuals in a population to estimate parasite prevalence. To estimate the total number of infected individuals, we multiplied the prevalence by the population size.

Next, we evaluated the size, or load, of individual infections across diet. Load is measured as the percentage of a worm’s total body area containing fluorescently-labeled parasites. For this measure, we were interested in the proliferation of infections inside of hosts, rather than initial exposures. We compared load only in infected worms (i.e., nematodes that had a fluorescent area greater than 0.001 μm^2^). It takes the parasite about two days to proliferate and generate new spores (Balla et al. 2016), so we subset the data to examine hosts that were at least two days old (L4 to young adult life stage). We determined through visual assessment of morphological traits (i.e. development of vulval window) that *C. elegans* with a body area greater than 20,000 μm^2^ were at least in the fourth larval or young adult stage. Then, we calculated median parasite load within that size class for each replicate population.

We first analyzed estimated population size as a function of diet, day, and their interaction. We fit a generalized linear model with a negative binomial error distribution. We then fit a generalized linear mixed model with a binomial error distribution to model a host’s infection status (yes/no) as a function of the days since exposure, diet, and their interaction. We included the population of origin (Plate ID) as a random effect to account for non-independence of hosts sampled from the same plate. Finally, we modeled parasite load in adult hosts, which was measured as the percentage of the host’s body area occupied by parasites. To account for plate-level variation, we took the median load of individuals measured in each population. We fit a generalized linear model in “glmmTMB” with a beta error distribution and analyzed the effect of diet, day of epidemic, and their interaction on this median load.

### General Information on Statistical Analyses

We performed all analyses in R Version 4.3 (R Core Team 2021) and made graphs using the “ggplot2” (Wickham 2016) and “patchwork” packages (Pedersen 2024). For linear models, we tested model assumptions using Levene (“car” package; Fox *et al*., 2018) and Shapiro-Wilk tests. For all generalized linear mixed models, we used the “DHARMa” package (Hartig 2022) to check that model assumptions were met. If the data failed to meet model assumptions for linear or generalized linear models, we used non-parametric tests. We tested the significance of our treatments using Type III tests with the Anova() function in the car package (Fox et al. 2018). For linear models, we performed F tests, and for generalized linear models, we performed Wald χ^2^ or Likelihood χ^2^ tests to assess the effect of our predictors. Then, we carried out post-hoc pairwise Tukey tests to compare between diets.

## Results

### Hosts feed less and acquire fewer parasites on low-quality diet

We estimated food uptake based on the accumulation of fluorescent beads. We found that feeding varied significantly across diets (Wald χ^2^; χ^2^ = 18.664, df = 2, p < 0.0001). In line with previous work (So et al. 2011b), we expected hosts on the low quality diet (DA837) to be smaller, which could itself cause hosts to acquire fewer beads in the assay. Confirming this, we found that larger hosts tended to eat more beads (Wald χ^2^; χ^2^ = 53.67, df = 1, p < 0.0001) and that host body size differed with diet (Kruskal-Wallis; χ^2^ = 7.71, df = 2, p = 0.02) (Figure S2). Hosts fed on the high quality diet (HB101) were 14% larger on average than those on the low quality diet (DA837; Pairwise Wilcoxon; W = 750, p = 0.018) and 10% larger than those on the medium quality diet (OP50; not significant; Pairwise Wilcoxon; W = 1309, p = 0.07) (Fig. S). We also found that bead ingestion increased with diet quality independently of host size: compared to hosts on the low quality diet, hosts of similar size on the medium quality diet consumed 2.85 times more beads (Tukey; z = −3.46, p = 0.002), and hosts of similar size on the high quality diet consumed about 5 times more beads (Tukey; z = −4.30, p < 0.0001) (Fig. 1A).

**Figure 1:**
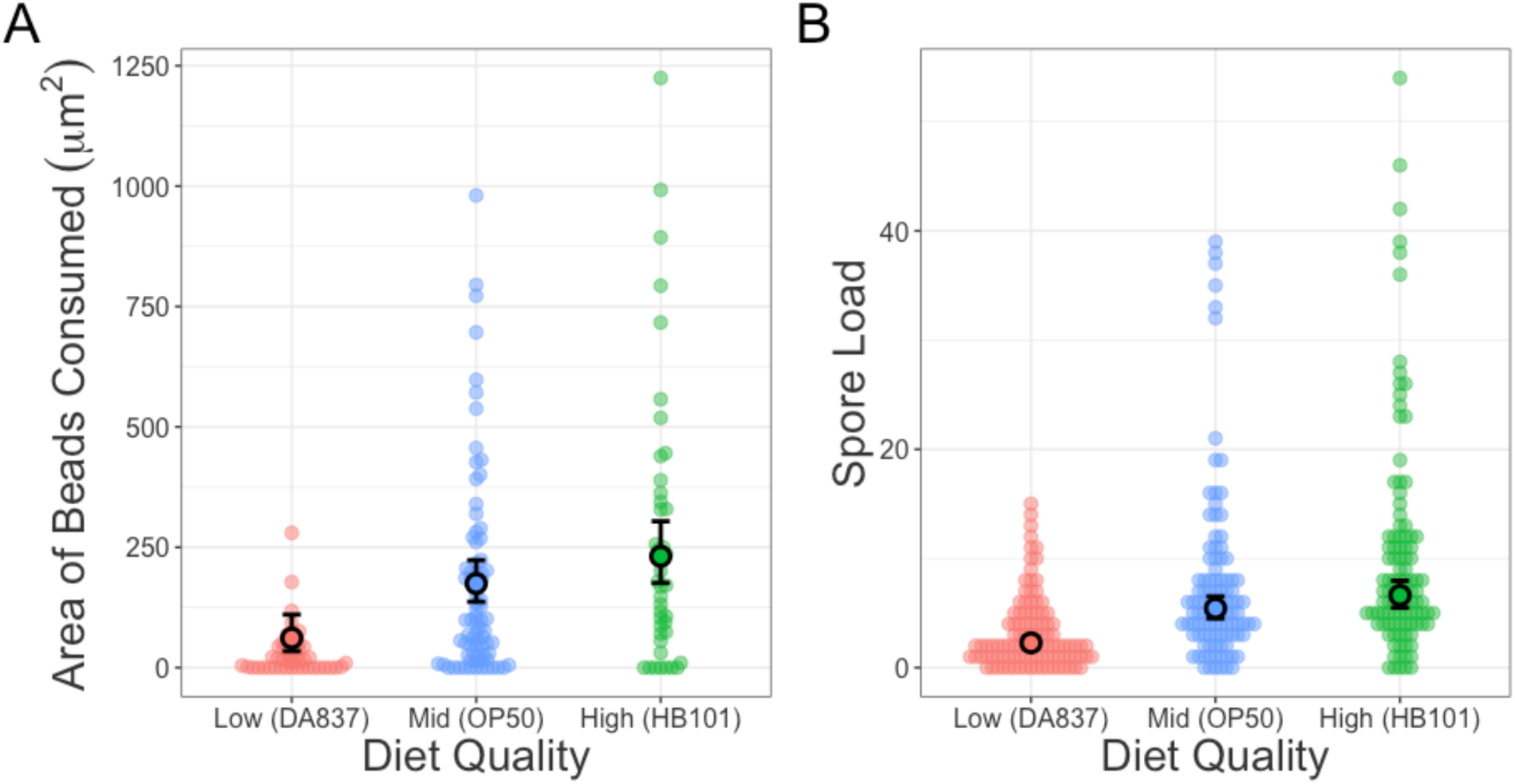
Hosts on low-quality diets eat less and acquire fewer spores. Transparent dots show raw values, and filled points indicate model-estimated marginal means for hosts of average size with 95% CI. A) Fluorescent bead acquisition, a proxy for food consumed, on each diet. B) Spore load after 24 hours of pathogen exposure.

Given that hosts on the low-quality diet ate less, we predicted that they would ingest fewer parasite spores. We found that parasite acquisition varied across diets in a 24-hour exposure (ANOVA; F_2, 312_ = 30.88, p < 0.0001). Hosts on the low-quality diet acquired the fewest parasites, with 17% of the sample having no established infections, as compared to 6% and 5% on medium- and high-quality diets respectively. Hosts on the high-quality diet had an estimated marginal mean of 6.62 sporoplasms (95% CI: [5.48, 7.95]), which was not significantly more than on the medium-quality diet, with 5.43 sporoplasms (95% CI: [4.50, 6.52]; Tukey; t = 1.46, p = 0.31). Hosts on the higher quality diets had significantly higher numbers of sporoplasms than hosts on the low-quality diet, with an average of 2.27 sporoplasms (95% CI: [1.80, 2.82]; Tukey for both comparisons; t = 7.27 and t = 6.08 respectively, p < 0.0001 for both) (Fig. 1B).

### Little evidence for differences in fitness costs of exposure across diets

Because hosts on the low-quality diet had fewer established sporoplasms, we expected reduced fitness effects of parasite exposure. To test this, we compared the fecundity of control and exposed hosts raised on the three diets. We found that exposure decreased lifetime fecundity by 27% on average (ANOVA; F_1, 117_ = 5.93, p = 0.016). Contrary to our prediction, hosts across diets had similar lifetime fecundity (ANOVA; F_2, 117_ = 1.58, p = 0.21) and similar reductions in fecundity with exposure (ANOVA Diet:Exposure; F_2, 117_ = 0.92, p = 0.40) (Fig 2A,B).

**Figure 2:**
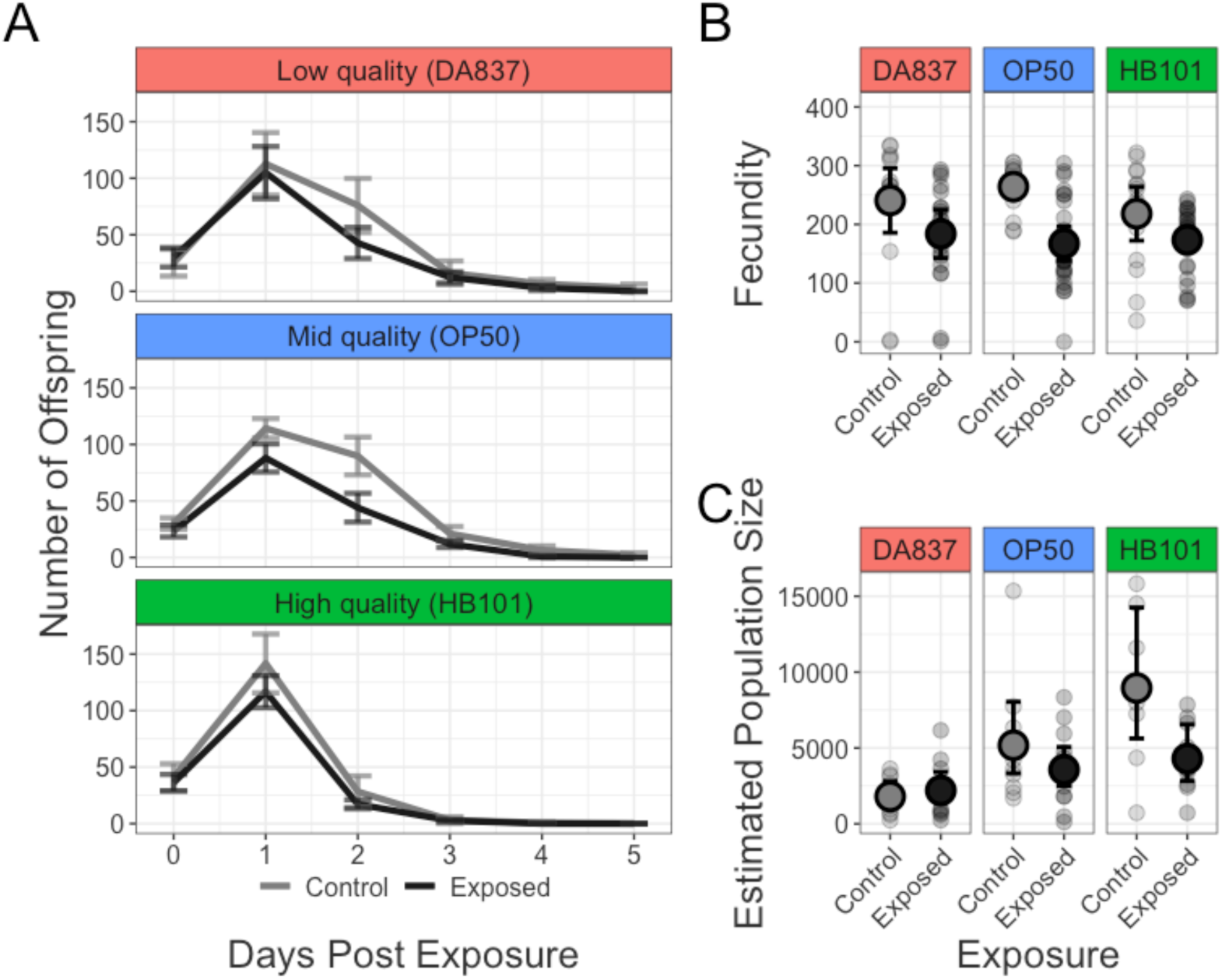
Disease impacts number and timing of offspring, regardless of diet. Transparent points show total number of offspring for individual hosts, and filled points represent the mean with 95% CI. A) The mean number of offspring produced each day in exposed and control conditions. B) Lifetime fecundity across diets. C) Sizes of control and exposed populations.

In line with previous results (Jiranek and Gibson 2023), reproductive timing was influenced by diet (Wald χ^2^ Diet:Day; χ^2^ = 78.08, df = 10, p < 0.0001) (Fig 2A). Peak fecundity occurred earlier for hosts on the high-quality diet, with exposed hosts producing 87% and control hosts producing 88% of their offspring over the course of the first two days of reproduction. In contrast, uninfected hosts on the lower quality diets produced only 57 and 55% of their offspring, respectively, over the same time frame. Exposed hosts tended to produce a greater proportion of their offspring on the first days of reproduction (Wald χ^2^ Day:Exposure; χ^2^ = 20.25, df = 5, p = 0.0013), likely because disease more strongly reduced fecundity on later days of reproduction as infections developed. Specifically, hosts exposed to *N. ironsii* on the lower quality diets produced a larger proportion of their total offspring (69 and 67%, respectively) in the first two days of reproduction relative to control hosts, while producing fewer offspring overall. Because hosts on the high-quality diet reproduced earlier, their fecundity schedule was relatively unaffected by exposure.

To determine whether differences in reproductive timing across diet and exposure affected population fitness, we measured population growth over multiple generations. Average population growth rate increased with diet quality (LR χ^2^; χ^2^ = 22.29, df = 2, p < 0.0001) (Fig. 2C). We found the largest populations on the high-quality diet, where the average size of uninfected populations was 8,955 (95% CI: [5,625, 14,257]). This was about 1.5 times the size of populations on the medium diet, which had a mean size of 5,178 (95% CI: [3,331, 8,049]), and nearly 5 times the size of populations on the low-quality diet, which had a mean size of 1,808 (95% CI: [1,163, 2,811]). There was no statistically significant effect of exposure (LR χ^2^; χ^2^ = 0.38, df = 1, p = 0.54) or interaction of exposure and diet (LR χ^2^; χ^2^ = 4.32, df = 2, p = 0.11). However, hosts that were exposed to the parasite on the high-quality diet had populations that were half the size of uninfected populations, with a mean size of 4,302 (95% CI: [2,824, 6,552]). On the medium quality diet, exposed populations were 31% smaller than in control conditions. In contrast, on the low-quality diet, exposed populations were 21% larger than control populations.

### Infection prevalence peaks faster on higher quality foods

We expected that the observed differences in parasite acquisition and establishment could affect the rate that infection spread through the population. To test this, we exposed populations of hosts on each diet to a low parasite dose and tracked population size and epidemic progression over four days. Supporting the results of our population growth assay, population sizes varied with diet (Likelihood ratio χ^2^; χ^2^ = 9.90, df = 2, p = 0.007). In general, host populations on the high-quality diet grew the fastest. By day 6 post exposure, the high quality diets had an estimated marginal mean population size of 287,202, nearly three-fold larger than that on the medium quality diet at 100,905 (Tukey; z = 6.13, p < 0.0001) and nearly twice that that on the low quality diet at 145,003 (Tukey; z = 4.01, p = 0.0002). In contrast to the population growth assay, population sizes on the low-quality diet overtook those on the medium-quality diet six days after exposure, but this difference was only marginally significant (Tukey; z = 2.03, p = 0.10) (Fig. 3A).

**Figure 3:**
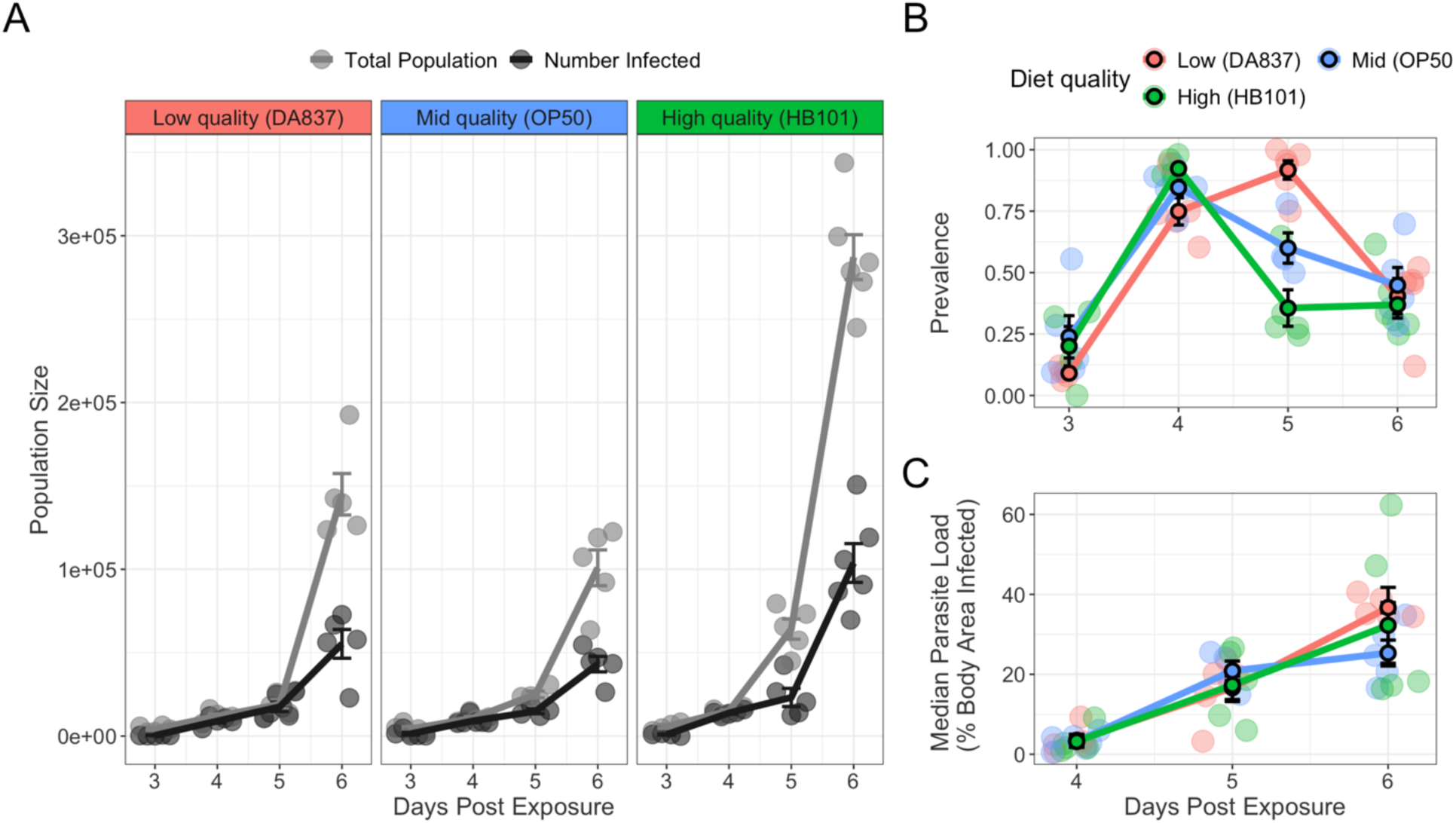
Diet influences magnitude and timing of epidemics. Transparent points are values for replicate populations, and error bars represent SEM of replicates within a treatment. A) Gray lines are population sizes, and black lines indicate the estimated number of infected individuals in these populations. B) Fraction of population with spreading infections in the gut. C) Median percentage of an adult host’s body with parasites (tagged with red fluorescence) for each replicate over time.

We then examined epidemiological dynamics. Overall, diet was a marginally significant predictor of disease prevalence (Wald χ^2^; χ^2^ = 5.09, df = 2, p = 0.08). Disease prevalence also varied depending on the time since parasite exposure (Wald χ^2^ Days post exposure; χ^2^ = 120.10, df = 3, p < 0.0001), and we detected an interaction of days post exposure with diet, indicating that the rank order of prevalence on each diet varied across the time series (Wald χ^2^ Days post exposure:Diet; χ^2^ = 66.47, df = 6, p < 0.0001). Consistent with the Parasite Acquisition assay, populations on the low quality diet had a lower prevalence of disease than populations on the medium (Tukey; z = 2.14, p = 0.08) and high (Tukey; z = 1.72, p = 0.20) quality diets early in the epidemic. Three days after exposure, only 9% of individuals on the low-quality diet were infected on average, compared to 24% on the medium and 20% on the high-quality diet.

Prevalence was significantly greater on HB101 than on DA837 four days post exposure (Tukey; z = 3.12, p = 0.005), with prevalence on OP50 falling in the middle. However, five days after exposure, populations on HB101 and OP50 declined in disease prevalence, with HB101 experiencing the sharpest decline to a mean prevalence of 34% (Tukey; DA837 (low) to HB101 (high): z = 7.28, p < 0.0001; DA837 (low) to OP50 (medium): z = 4.60, p < 0.0001; HB101 (high) to OP50 (medium): z = 2.43, p = 0.04). This decline coincided with large population expansions, suggesting that while the total number of infected individuals continued to increase (Fig. 3A), the population contained many newborn hosts that had not yet acquired infection. In contrast, prevalence continued to increase on the low-quality diet on the fifth day post exposure, climbing to a mean of 91%. By the sixth day post exposure, populations on the low-quality diet experienced a dip in prevalence, leading to relatively equal prevalence across all diets. Despite having similar prevalence, populations on different diets had large differences in the numbers of infected individuals. By the sixth day post exposure, there were an estimated average of 103,791 infected hosts in populations on the high-quality diet, compared to 43,163 on the medium quality and 55,301 on the low quality (Fig. 3B).

To assess whether diet affected the progression of host infection, we measured the parasite load of adult hosts. We calculated the percentage body area occupied by parasites. We found that mean parasite load increased throughout the experiment (Wald χ^2^; χ^2^ = 41.82, df = 2, p < 0.0001), but that diet did not affect parasite load (Wald χ^2^; χ^2^ = 0.005, df = 2, p = 0.99) (Fig. 3C). Load was relatively low in adults after four days of exposure, with a mean body area occupied by parasites of about 3-4% across diets. By five days post exposure, overall load in infected adults was higher, with about 17% of body area consumed by parasites for hosts on the low- and high-quality diets and 21% for hosts on the medium quality diet. By Day 7, hosts raised on the low-quality diet had slightly higher parasite loads (mean: 37%) compared to the medium quality diet (mean: 25%), but this was not statistically significant. The high-quality diet was intermediate between the two at 32% parasite load.

## Discussion

Host diet shapes epidemic progression through interconnected mechanisms. Using a study system that allows detailed individual and population-level measures of disease, we tested ways in which diet can alter the transmission of parasites, thereby altering epidemics. We tested the role of diet in parasite acquisition, host fitness, and population density in a series of experiments on three host diets that varied in quality. We found that diet affected both parasite acquisition and population growth, but found little evidence that diet altered within-host infection dynamics. Epidemics progressed more quickly in host populations on high-quality food, which we attribute to increased host density and parasite acquisition.

### Diet alters feeding and parasite acquisition

We predicted that diet quality would drive host feeding behavior, thus leading to different rates of parasite acquisition. As seen in other host–parasite systems (Penczykowski et al. 2014, Davenport et al. 2025), hosts increased feeding rate as dietary quality increased, thereby encountering more parasite spores (Fig. 1). We found that hosts on the higher quality diets were more likely to have a least one germinated sporoplasm and had a higher number of total sporoplasms. Perhaps due to this greater parasite acquisition on the higher quality diets, disease prevalence increased faster on these diets in the early days of the epidemic experiment (Fig. 3B). Host populations on the low-quality diet experienced peak disease prevalence a day later, suggesting that lower parasite acquisition delayed the epidemics on this diet.

A delay of one day is epidemiologically relevant in this system. *C. elegans* has a rapid generation time (∼4 days) and progresses rapidly through developmental stages (Félix and Braendle 2010). Thus, a delay of a single day could make the difference between getting infected in the fourth larval stage (sub-adult) rather than the first larval stage (newborn). For an individual host, infections acquired at the first larval stage have much more severe fitness consequences than infections acquired one day later, at the fourth larval stage (Balla et al. 2015). This might explain why populations on the worst quality food grew larger than populations on the medium quality food during the epidemic experiment (Fig. 3A) despite growing slower in the population growth experiment (Fig. 2C): a greater number of hosts may have avoided early infection and contracted disease later in life when infection had less impact on fitness. Altogether, we find that diet modulates a host’s risk of contacting parasite spores, such that hosts that ate less on low quality foods had a lower risk of infection.

### No evidence for diet impacts on within-host infection and fitness effects

Parasite load can affect fitness consequences of disease and epidemiological dynamics (Anderson and May 1982). Given that hosts on low-quality diets ingested fewer parasites (Fig 1B), we predicted that they would have lower burdens of disease overall. Instead, we did not detect differences in disease burden in infected adult hosts in an epidemic context (Figure 3C). This result contrasts with a review that found increasing diet quality was typically found to increase parasite load in invertebrate hosts (Cressler et al. 2014). For example, high quality food increased the viability of parasite spores in their zooplankton hosts (Penczykowski *et al*., 2014; Narr *et al*., 2019), and nutrient supplementation exacerbated fungal and bacterial disease in corals (Bruno et al. 2003).

There are several reasons that higher rate of parasite acquisition and establishment might not lead to higher disease burdens. First, the parasite is likely virulent even in smaller doses: infections deriving from a single spore can grow to encompass the whole gut in a closely-related *C. elegans–Nematocida* interaction (Balla et al. 2016). Alternatively, host resistance to disease is frequently greater for hosts raised in high quality environments (Moret and Schmid-Hempel 2000; Kraaijeveld & Godfray, 1997; Fellous and Lazzaro 2010). Thus, increased resistance of nematode hosts raised on the higher quality diet may have prevented parasites from spreading as quickly, tempering the higher rate of parasite acquisition.

Fecundity effects of parasite exposure also did not differ with diet. Hosts on the lowest quality diet suffered reproductive losses to disease of a similar magnitude to the other diets in spite of ingesting fewer spores (Figure 2B). Similarly, hosts raised on the high-quality diet, which ingested the greatest number of parasites, did not suffer greater fecundity reductions than hosts on the other diets. However, hosts on the high-quality HB101 diet showed the largest reduction in population size with exposure during the Population Growth assay (Fig. 2C). It is possible that the faster development of hosts raised on HB101 allowed hosts to experience more generations, thus widening the gap in population size between healthy and exposed populations (Jiranek and Gibson 2023). Alternatively, *N. ironsii* virulence might be greater in a high-density environment. Theory suggests that resource supply can generate variation in parasite virulence (Lively 2006, Hall et al. 2009b), and empirical work has shown that high host density increases parasite virulence (Lively et al. 1995, Bell et al. 2006). Increased density on the high-quality diet could have increased resource scarcity among hosts, thus leading to increased virulence of disease.

Overall, our study detected relatively little contribution of within-host dynamics to epidemic processes. Although the low-quality diet had a later peak in disease prevalence in the epidemic, infected hosts had a similar parasite burden. Thus, greater parasite acquisition on high quality diets likely modulated epidemics through increasing the number of individuals that contacted the parasite rather than the number of spores per host. We did, however, conduct this work on a single host strain, N2, that has relatively low resistance. Previous work suggests that there is genetic variation in resistance among host strains, with some able to clear infections from the gut (Balla et al. 2015). It may be that the effect of diet is larger in cases where hosts have a more active resistance defenses, so future studies should investigate how this interaction between host strain and diet drives epidemics.

### Diet alters epidemics through impacts on host density

Perhaps the largest impacts on the course of epidemics occurred because of diet-dependent effects on host population growth. Theory and empirical work support faster spread of disease in the larger, denser populations supported on high quality diets (Anderson and May 1981). Hosts on the high-quality HB101 grew (So *et al*., 2011a;) and reproduced (Fig. 2A) faster, which accelerated population growth (Fig. 2C). Because *C. elegans* development involves achieving certain “threshold” sizes to molt, a faster growth rate leads to faster progression through molts and achievement of reproductive maturity (Uppaluri and Brangwynne 2015).

Accelerated development and population growth on the high-quality diet led to significant changes to disease dynamics: when hosts on the high-quality diet exhibited a burst of reproduction, disease prevalence dropped rapidly with the influx of new uninfected juveniles.

Thus, although the absolute number of infected individuals continued to grow, the proportion of infected hosts in the population was lower (Fig. 3A,B). This “safety in numbers” was also seen in a snail host with its trematode parasite; in populations that were manipulated to have high densities, total recruitment of a trematode parasite increased, but per-capita infection risk for the hosts decreased (Buck et al. 2017). Our study provides a proof-of-concept that increased host density supported on high-quality diets can cascade to influence epidemiological dynamics.

### Conclusion

Supporting previous work, we find that diet quality affects feeding behavior and reproduction. We extend upon this work by demonstrating that these individual-level changes scale to affect disease spread at the population level across replicated experimental epidemics.

In contrast to other studies, we find relatively little evidence that epidemic size is driven by differences in within-host disease spread. Through its impacts on contact rate with trophically-acquired parasites and host population density, diet determines the size and timing of epidemics.

## Acknowledgments

*E. coli* strains were provided by the CGC, which is funded by the NIH Office of Research Infrastructure Programs (P40 OD010440). The host line (*C. elegans* N2) and parasite strain (*N. ironsii*) were obtained from the Troemel lab at UCSD. The authors thank Anne Janisch, Sarah Hesse, Irish Amundson, Caroline Amoroso, and Leila Shepard for helping with experiments and the Gibson and Galloway labs for helpful comments on the manuscript. This work was supported by supported by funding from the National Institute of General Medical Sciences (R35 GM137975-01) and the Jeffress Trust Awards Program in Interdisciplinary Research. JAJ was also supported by an NSF Research Training Grant (2021791).

## Conflict of interest statement

The authors declare no conflict of interest.

## Notes

### Competing Interest Statement

The authors have declared no competing interest.

