## Supplementary Information for "Diet alters epidemic size and timing in a trophically-transmitted parasite"

Supplementary Material for **Diet alters epidemic size and timing in a trophically-transmitted parasite**

**Figure S1: Conceptual overview of questions and assays included.**

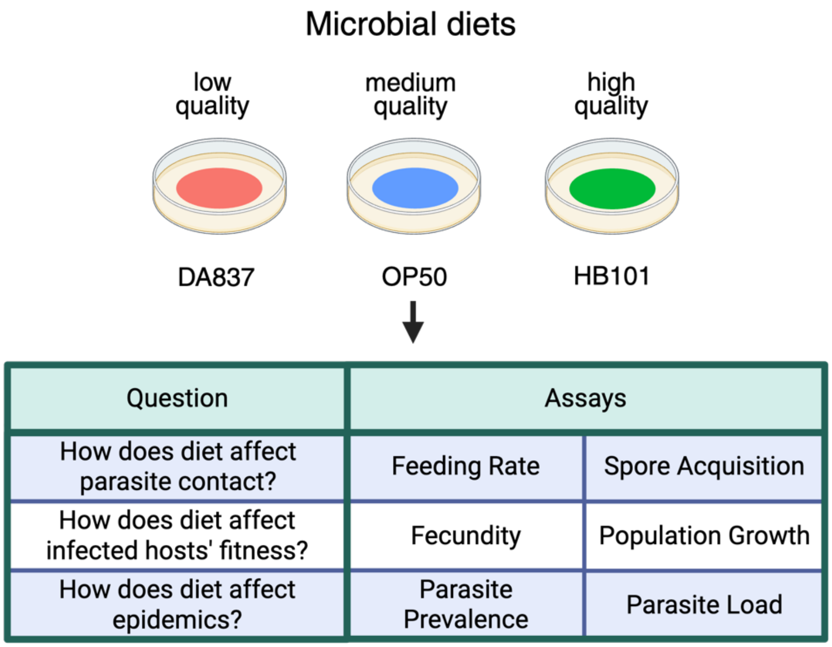


**Figure S2: Hosts grow larger on the highest quality diet.** Transparent dots show raw values of host size measured from a 2D microscope image, and error bars present means and normal 95% confidence intervals.

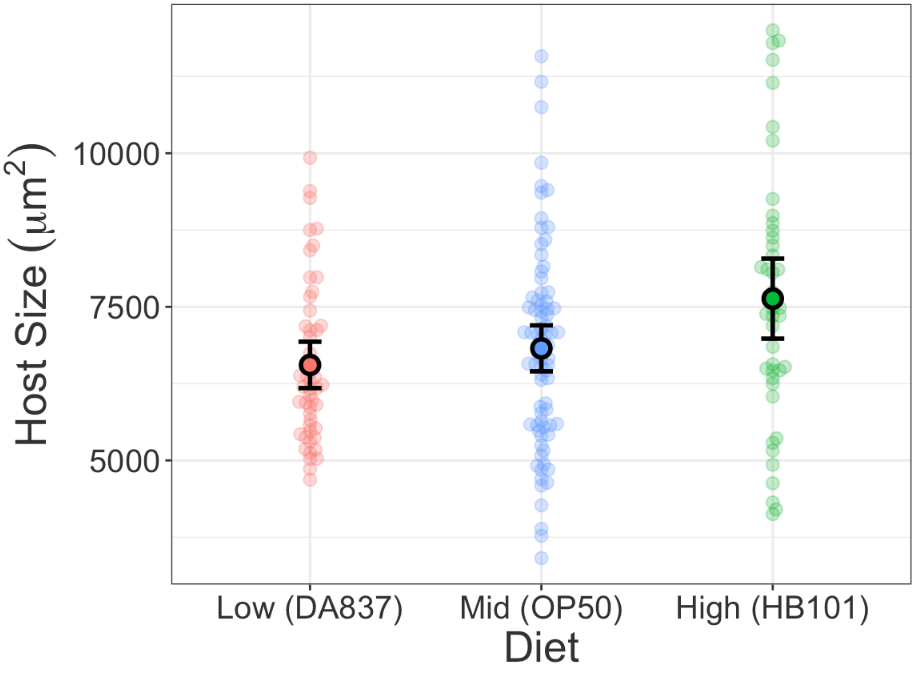

**Figure S3: Feeding rate does not differ between control and parasite-exposed hosts.** Transparent dots show raw values for fluorescent bead acquisition, a proxy for food consumed, on the OP50 diet. Filled points indicate model-estimated marginal means with 95% CI.

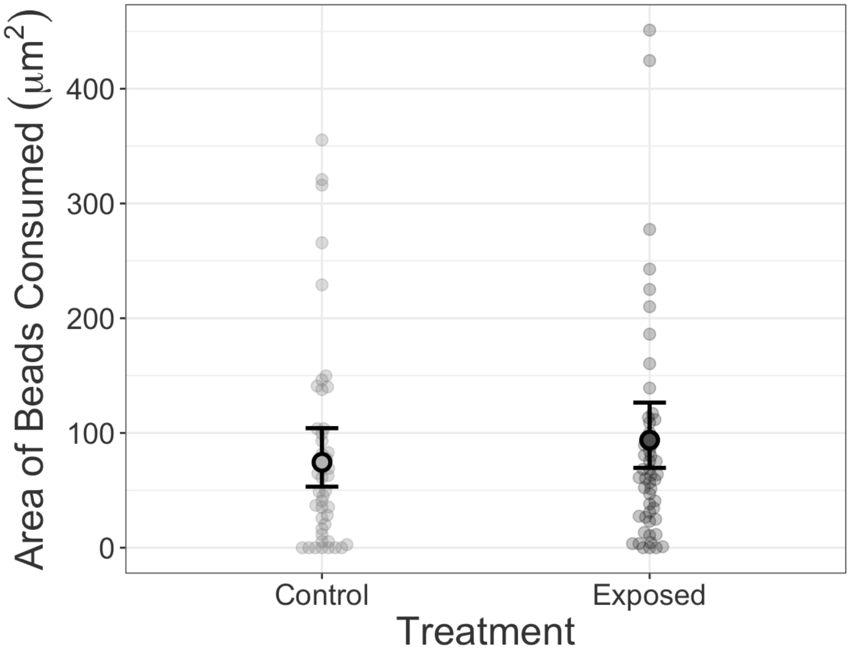


**Appendix A: Feeding in Exposed Hosts**

In some systems, parasites can affect host feeding rate (Hite et al. 2020). Previous experiments measuring pharyngeal pumping to measure feeding rate suggest that *C. elegans* cannot detect *Nematocida* spores and do not change their feeding rate when spores are present in food (Troemel et al. 2008). To test this using our fluorescent bead assay approach, we compared feeding rates of hosts exposed to *N. ironsii* spores to that of hosts that were not exposed to the parasite.

Hosts were reared to the L1 stage on the medium quality OP50 diet. Then, 1,000 hosts were moved to either control or exposed agar plates (n = 3 plates per treatment). As in the unexposed Bead Acquisition Assay, these assay plates had approximately $1.27\times10^{10}$ 0.10 micron fluorescent polystyrene beads mixed in with the food. Control plates had OP50 bacteria treated with 120 μl of control lysate and exposed plates had 120 μl of lysate containing approximately $8.90\times10^{6}$ *N. ironsii* spores. This was twice the dose we used in the Parasite Acquisition assay because in our Bead Acquisition Assay, we used twice the number of hosts. To keep the ratio of spores per host constant while also maintaining the larger populations needed for this assay, we doubled the dose.

After 30 minutes of foraging, we washed hosts off the assay plates, anesthetized and fixed them on microscope slides, and photographed individual hosts using a fluorescent microscope as described in the Bead Acquisition Assay (Control: n = 45; Exposed; n = 50). The images were analyzed in ImageJ by a researcher who did not know the treatment of each photograph. The data had a negative binomial error distribution, so we analyzed it in a model implemented with "glmmTMB" that included parasite exposure as a predictor and the assay plate as a random effect.

We found no evidence that parasite exposure affected feeding rate (Wald $\chi^2$; $\chi^2 = 1.45$, df = 1, p = 0.23). Hosts in the control condition ingested a mean of 74.4 $\mu m^2$ beads (95% CI: [53.2, 104]), whereas hosts in the exposed group ingested an average of 93.8 $\mu m^2$ beads (95% CI: [69.5, 126]). In contrast to work in other systems, we do not detect behavioral avoidance of food contaminated with parasites (Gibson and Amoroso 2022).

**Table S1: Summary of replication, sample size, and censored data for assays**

**A. Bead acquisition**

| Diet | Replicate Plate[1] | Hosts measured[2] |
|---|---|---|
| Low quality (DA837) | 1 | 12 |
| | 2 | 3 |
| | 3 | 8 |
| | 4 | 10 |
| | 5 | 1 |
| | 6 | 9 |
| | 7 | 6 |
| Mid quality (OP50) | 1 | 4 |
| | 2 | 7 |
| | 3 | 8 |
| | 4 | 8 |
| | 5 | 19 |
| | 6 | 7 |
| | 7 | 15 |
| | 8 | 10 |
| High quality (HB101) | 1 | 5 |
| | 2 | 6 |
| | 3 | 4 |
| | 4 | 7 |
| | 5 | 13 |
| | 6 | 6 |
| | 7 | 2 |

[1]Number of independent agar plates that hosts fed on during assays

[2]Number of hosts fixed, photographed, and scored

**B. Bead acquisition: exposed hosts**

| Treatment | Replicate Plate | Hosts measured[1] |
|---|---|---|
| Control | 1 | 18 |
| | 2 | 16 |
| | 3 | 11 |
| Exposed | 1 | 9 |
| | 2 | 29 |
| | 3 | 12 |

**C. Parasite acquisition**

| Diet | Hosts measured |
|---|---|
| Low quality (DA837) | 106 |
| Mid quality (OP50) | 105 |
| High quality (HB101) | 105 |

[1]Number of hosts fixed, photographed, and scored

**D. Fecundity**

| Block | Diet | Exposure | Host replicates | Number censored |
|---|---|---|---|---|
| Block 1 | Low quality (DA837) | Control | 7 | 0 |
| | | Exposed | 12 | 2 |
| | Mid quality (OP50) | Control | 15 | 0 |
| | | Exposed | 25 | 2 |
| | High quality (HB101) | Control | 15 | 2 |
| | | Exposed | 25 | 2 |
| Block 2 | Low quality (DA837) | Control | 10 | 1 |
| | | Exposed | 10 | 0 |
| | Mid quality (OP50) | Control | 5 | 0 |
| | | Exposed | 5 | 0 |
| | High quality (HB101) | Control | 5 | 0 |
| | | Exposed | 5 | 0 |

**E. Population growth**

| Diet | Exposure | Replicate populations | Number censored |
|---|---|---|---|
| Low quality (DA837) | Control | 55 | 1 |
| | Exposed | 55 | 0 |
| Mid quality (OP50) | Control | 60 | 2 |
| | Exposed | 65 | 1 |
| High quality (HB101) | Control | 40 | 1 |
| | Exposed | 55 | 0 |

**F. Epidemic progression**

| Diet | Day | Replicate populations | Censored |
|---|---|---|---|
| Low quality (DA837) | 3 | 5 | 0 |
| | 4 | 5 | 0 |
| | 5 | 6 | 0 |
| | 6 | 5 | 0 |
| Mid quality (OP50) | 3 | 5 | 0 |
| | 4 | 6 | 0 |
| | 5 | 4 | 1 |
| | 6 | 5 | 0 |
| High quality (HB101) | 3 | 4 | 1 |
| | 4 | 5 | 0 |
| | 5 | 5 | 0 |
| | 6 | 6 | 0 |
